# Automatic Reconstruction of Cerebellar Cortex from Standard MRI Using Diffeomorphic Registration of a High-Resolution Template (ARCUS)

**DOI:** 10.1101/2020.11.30.405522

**Authors:** John G. Samuelsson, Bruce Rosen, Matti S. Hämäläinen

**Affiliations:** Massachusetts General Hospital - Massachusetts Institute of Technology - Harvard Medical School; Harvard-MIT Division of Health Sciences and Technology, Athinoula A. Martinos Center for Biomedical Imaging, 149 13th St., Charlestown, MA 02129, USA

## Abstract

As cumulating evidence points to a wider range of functional tasks and neurological conditions that involve the cerebellum than previously known, the interest for examining the cerebellum with non-invasive neuroimaging techniques is growing. However, the standard methods of computational neuroanatomy for segmenting and reconstructing the cerebral cortex work poorly for the cerebellar cortex at the resolutions attainable with contemporary MRI technology because of its extremely intricate folding, making detailed and topologically correct reconstructions of the geometry of the cerebellar cortical surface unfeasible. Recently, a detailed surface reconstruction of the human cerebellar cortex was achieved from an ex-vivo specimen. These novel anatomical data enable a new reconstruction technique where this detailed surface reconstruction is morphed to subject space based on standard in-vivo MRI data. The result is an approximate reconstruction of the cerebellar cortex that requires only standard-resolution MRI data and can be used e.g., in functional neuroimaging, for integrating topographic population data or for visualizing topographic data on flattened surface patches.

## Introduction

The cerebellum has historically been assumed to relate mainly to lower-level functions such as posture and motor coordination. The tentorium cerebelli that separates the cerebellum from the occipital lobe has even been compared to a “Maginot Line” that separates the higher-order functions of the cerebral cortex with the lower-order functions of the cerebellum (Andreasen and Pierson, 2008). With the advent of new neuroimaging technologies, this idea has been challenged and a new view has emerged where the cerebellum is central in a wide range of functional tasks including various aspects of cognitive processing such as working memory and language (Middleton and Strick, 2001; Schmahmann and Sherman, 1998; Stoodley and Schmahmann, 2009). Lending credence to this idea, it is now well-known that the cerebellum contains over 70 % of the brain’s neurons and is densely interconnected with both the sub- and neocortex (Andersen et al., 1992; Herculano-Houzel, 2009). Furthermore, it was recently found that the areal ratio of the cerebellar to the cerebral cortex in humans is 80 %, while this number in Macaques is just 35 %, implying that the cerebellar cortex has expanded more than the neocortex in the evolutionary step between non-human and human primate, suggesting a cerebellar role in distinctively human traits, e.g., language (Sereno et al., 2020). From a computational perspective, it has also been suggested to be the supervised learning central of the human brain (Albus, 1971; Marr, 1969; Raymond and Medina, 2018).

Along with our advancing understanding of the wide range of roles that the cerebellum plays in health and disease, it becomes increasingly more pressing to having access to adequate tools with which to examine cerebellar functional neurophysiology. Recent studies have suggested that cerebellar electrophysiology can be measured non-invasively with magneto- and electroencephalography (MEG and EEG; Andersen et al., 2020; Samuelsson et al., 2020b). However, reconstructing the neural currents based on M/EEG sensor data, the inverse problem, is a formidable challenge that is considerably alleviated by having access to a surface source space which utilizes prior anatomical knowledge (Hillebrand and Barnes, 2003; Okada, 1989). A surface source space is also helpful for visualization purposes. Currently, however, there is no way of reconstructing the cerebellar cortex adequately from in-vivo MRI data because the resolution is insufficient. A resolution of < 0.2 mm isotropic voxel size is required to properly segment and reconstruct the cerebellar cortex, while contemporary MRI voxel resolution is usually 1 mm, or 0.75 mm in newer state-of-the-art systems (Sereno et al., 2020). We are thus still far away from achieving sufficient resolution to reconstruct the cerebellar cortex using conventional neuroanatomical segmentation methods.

Previous cerebellar atlas methods have focused on classification of voxels into anatomical or functional regions (Buckner et al., 2011; Diedrichsen et al., 2009; Jenkinson et al., 2012; Makris et al., 2003; Ren et al., 2019). In some cases these voxels have been projected onto flattened surface patches (flat maps), which enables visualization of topographic data on a flat surface (Diedrichsen and Zotow, 2015; Makris et al., 2005). Because of the unfeasibility to properly reconstruct the cerebellar cortex from standard MRI data, however, these flat maps are based on standard gray/white matter segmentations of the cerebellum which capture little more than its convex hull, leading to poor flat map representations of the actual cortex. While these atlases and segmentation tools are excellent for categorizing volumetric data into structural or functional regions and defining the outer boundaries of the cerebellum and its regions, and we indeed make use of these tools in the current study, they do not represent veridical reconstructions of the cerebellar cortex. A technique for proper reconstruction and flat map representation of the cerebellar cortex, and parcellation of this cortex, is thus still missing. The present study aims to address this missing link.

An alternate method that does not rely on the traditional segmentation-reconstruction approach is to use a diffeomorphic technique where another reconstruction is registered to the subject. It is essentially the inverse of the morphing from a subject to a common space, e.g., Talairach or MNI, that is currently being done when integrating neuroimaging population data. The difference here is that the common space is a high-resolution reconstruction of the cerebellar cortex that is otherwise unfeasible to achieve. This technique is made possible by the recent full reconstruction of the cerebellar cortex from ex-vivo scans by Sereno et al. (2020). This manuscript presents such a method, which we call ARCUS (Automatic Reconstruction of Cerebellar cortex from standard MRI USing diffeomorphic registration of a high-resolution template). We test the method on standard resolution (1 mm) MRI data. Although a full validation would require comparisons to multiple reconstructed surfaces which are unavailable at present, we compared the reconstruction to a high-resolution (0.25 mm) data set, which gives a sense of the accuracy of the reconstruction.

ARCUS is fully automatic and requires only a FreeSurfer reconstruction data set that includes the Aseg segmentation along with the necessary Python code and its dependencies.

## Methods

This study and the code for implementing this technique was done in Python 3.7, relying on the software packages Nibabel version 3.1.0 for handling neuroimaging data, Evaler for handling geometric data and ANTs version 0.2.5 for performing the non-linear registration (Avants et al., 2011; Brett et al., 2020; Samuelsson et al., 2020a). The workflow of ARCUS is illustrated in **Figure 1** and described in the following steps:

1. First, the part of the subject’s MRI volume that contains the cerebellum is isolated using FreeSurfer’s automatic segmentation of subcortical structures (Aseg) (Fischl, 2012). With Aseg, the cerebellum is further subdivided into a rough white matter and cortex segmentation which is used to compute the average contrast values of the white matter and cortex voxels.
2. A high-resolution reconstruction of the cerebellar cortex from an ex-vivo human cerebellum, presented in detail in (Sereno et al., 2020), is then loaded and fitted to the cerebellum by means of an affine transformation.
3. The reconstructed cerebellar cortex, which is a surface manifold containing 4.6 million vertices and 9.2 million triangular faces, is transformed into a “fake” MRI volume by assigning the average contrast value of the cerebellar white matter to all voxels inside the surface and assigning the contrast value of the cerebellar cortex to all voxels that contain any piece of the surface manifold. This artificial MRI volume is then resampled to fit the size of the subject data and all vertices in the manifold are rescaled and translated to fit the resampled MRI volume. The subject MRI volume is then thresholded such that all voxels whose contrast is less than 10% higher than the average cortical voxel is put to zero. This is to better distinguish the lobules, particularly in the anterior lobe which tend to blur together. The artificial MRI volume is then registered to the subject MRI using the non-linear deformation procedure symmetric normalization with cross-correlation as the optimization metric as implemented in the ANTs software package.
4. The velocity fields from this transformation are saved and then applied to the vertices in the surface manifold, thus registering the surface manifold to the subject volume. The non-linear registration procedure is then repeated but this time without thresholding.
5. Finally, the registered surface is put pack into the original subject MRI coordinate system and vertex normals in the fitted surface mesh are recalculated.

**Figure 1.**
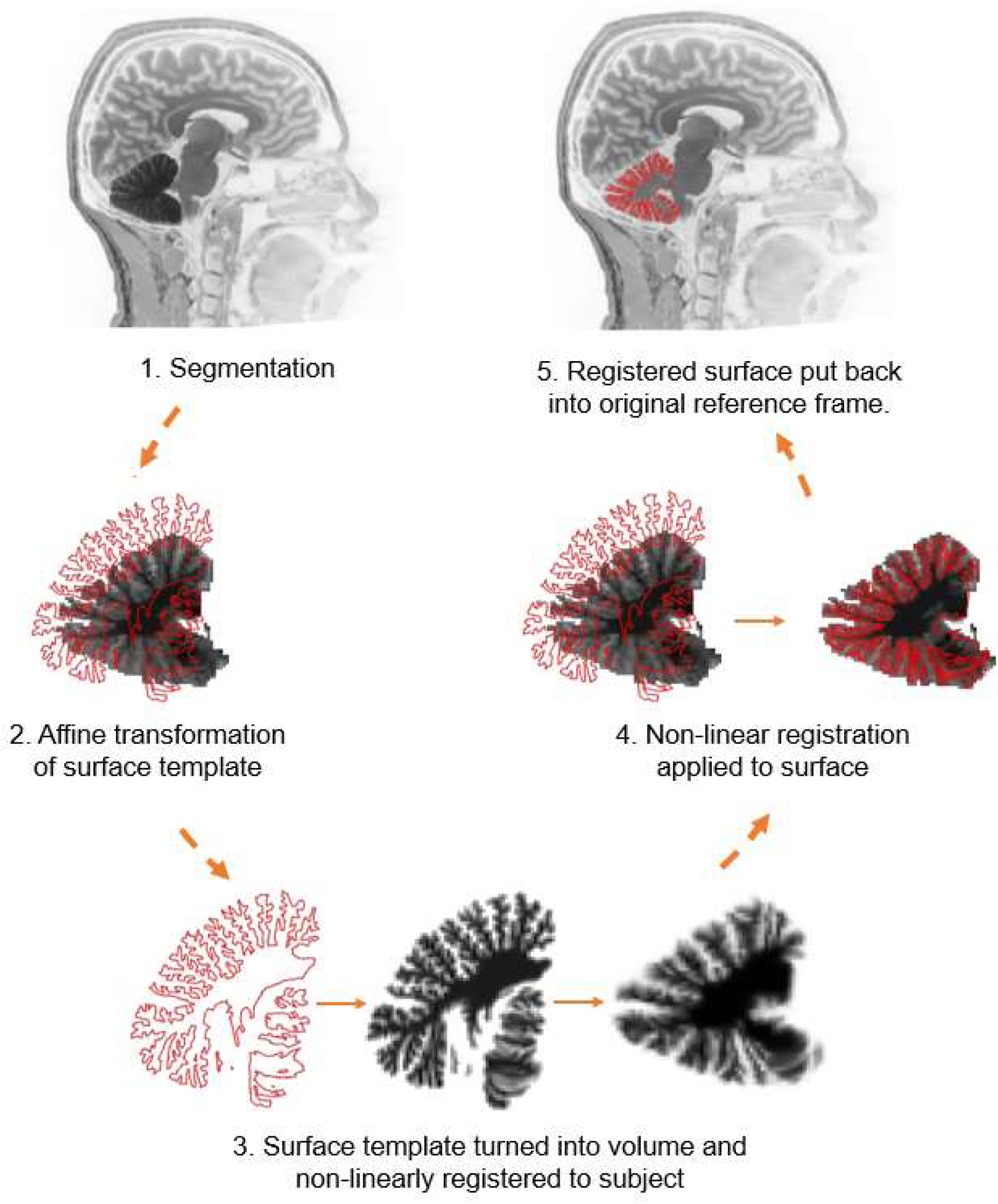
Conceptual schematic of reconstruction method outlining the major steps. 1. Cerebellum is isolated and white matter and cortical voxels are roughly segmented using FreeSurfer Aseg. 2. High-resolution surface template (red) is loaded and approximately fitted to subject cerebellum by means of an affine transformation. 3. Template surface is turned into an artificial MRI volume using the average contrasts from the FreeSurfer Aseg segmentation and volumetric template is registered to the subject MRI using ANTs non-linear SyNCC registration. 4. The velocity fields of the registration are applied to the vertices in the template surface, registering the surface to the subject’s cerebellum. 5. The registered surface is put back into the subject’s original reference frame.

The computational time cost of the reconstruction varies depending on the resolution; a higher resolution will give a better result but increase the time complexity. For the standard 1 mm^3^ resolution, the reconstruction takes about two minutes using a single core on an Intel Xeon Gold 6130 Processor with a base frequency of 2.10 GHz. The memory storage of the pre-processed template data required for the reconstruction is 1.7 Gb.

In Sereno et al. (2020), the high-resolution surface was divided up into different anatomical regions which were separately flattened. The cortex was also inflated. Because there is a one-to-one mapping between the vertices in the flat map, the inflated cortex and the surface manifold, which can be fitted to a subject, subject data can be projected to and visualized in the flat map and the inflated cortex. This is equivalent to a morphing procedure to a common space. **Figure 2** shows the cortical parcellation over the cerebellar cortex that has been fitted to a subject, also visualized on the inflated cortex and the flat map, which has been assembled from the flattened patches in Sereno et al. (2020). The cortex shown in Figure 2 is a downsampled version of the original tessellation with 200000 faces, which is easier to handle in computations.

**Figure 2.**
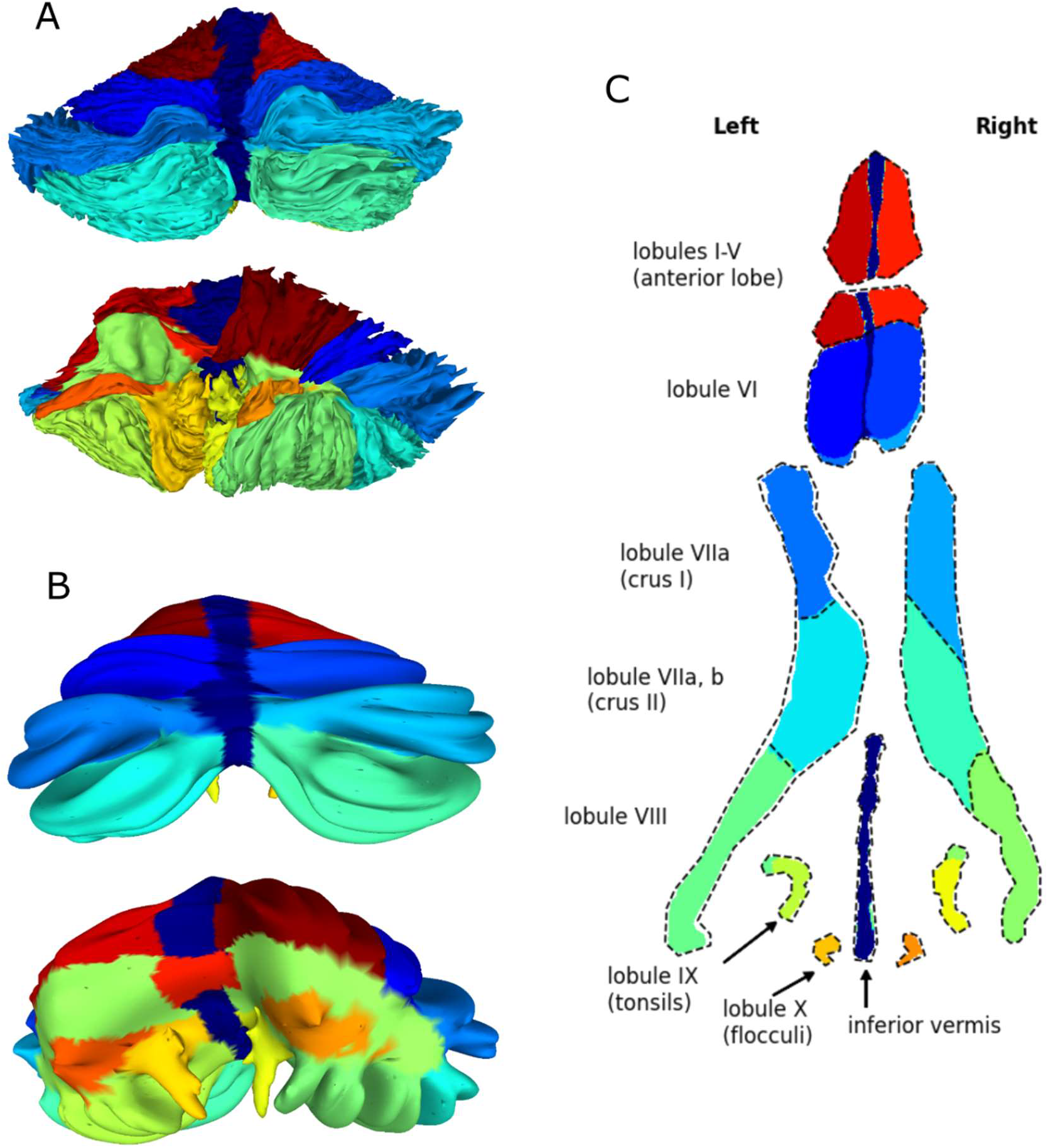
Parcellation of cerebellar cortex visualized on a cerebellar cortical reconstruction (A), inflated cortex (B) and flat map (C). The high-resolution (0.25 mm isotropic) MRI data that the technique was tested on was retrieved from Lüsebrink et al. (2017) and the standard-resolution (1 mm isotropic) MRI data was from Samuelsson et al. (2020a).

## Results

Results based on the standard resolution (1 mm isotropic) MRI data are shown in **Figure 3** which displays 3D reconstructions done with ARCUS and Aseg as well as the intersections of sagittal and coronal cross-sections in the intermediate zone with the cortical reconstructions. The reconstruction generates a largely correct convex hull of the cerebellum and the overall lobular-level cortical reconstructions are satisfactory in the posterior potions between the major cerebellar fissures. The performance is worse in the anterior lobe, however, where the fissures separating the lobules are smaller and the reconstructed lobules sometimes ending up traversing the smaller fissures.

**Figure 3.**
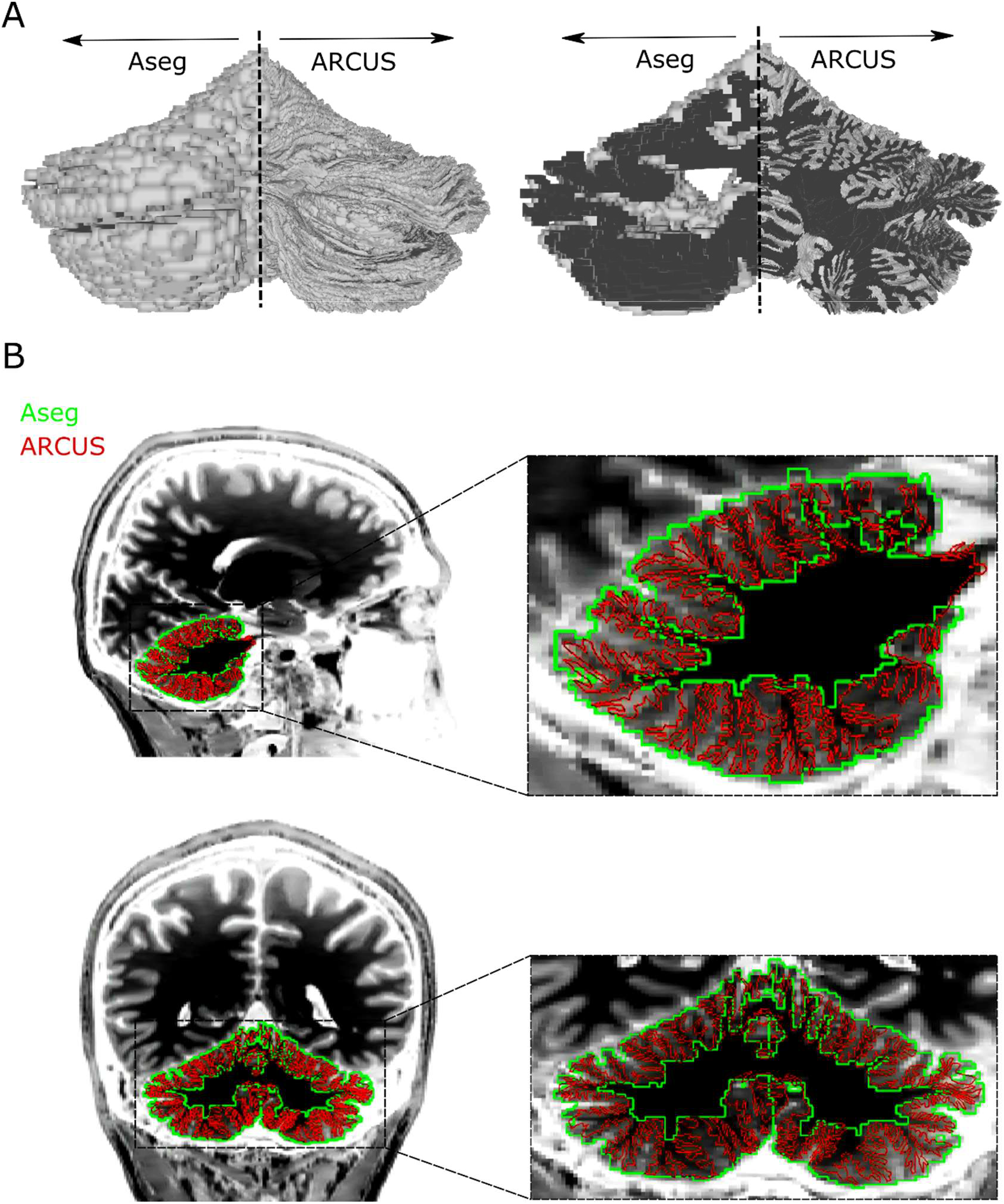
Reconstruction of the cerebellar cortex from standard (1 mm isotropic) resolution MRI data. The reconstruction’s intersection with the cross-section is marked in red, superposed on the original MRI data and the FreeSurfer tessellation of the cerebellar cortex is marked in green. A) 3D reconstructions of the cerebellar cortex using Aseg and ARCUS, respectively. B) Sagittal and coronal cross-sections of a subject MRI together with the reconstructed cerebellar cortex and the FreeSurfer Aseg tessellation.

**Figure 4** shows the reconstruction together with the high-resolution MRI data. These data have been downsampled to standard 1 mm resolution prior to performing the reconstruction. To better illustrate the quality of the reconstruction, it is superposed on the high-resolution volumetric data. Again, we note that the quality of the reconstruction is generally better in the posterior part of the cerebellum but worse in the anterior region, as well as in the most lateral portions. With the higher resolution data, we can also note that the lobules are reconstructed well but the smaller folia are often not adequately captured.

**Figure 4.**
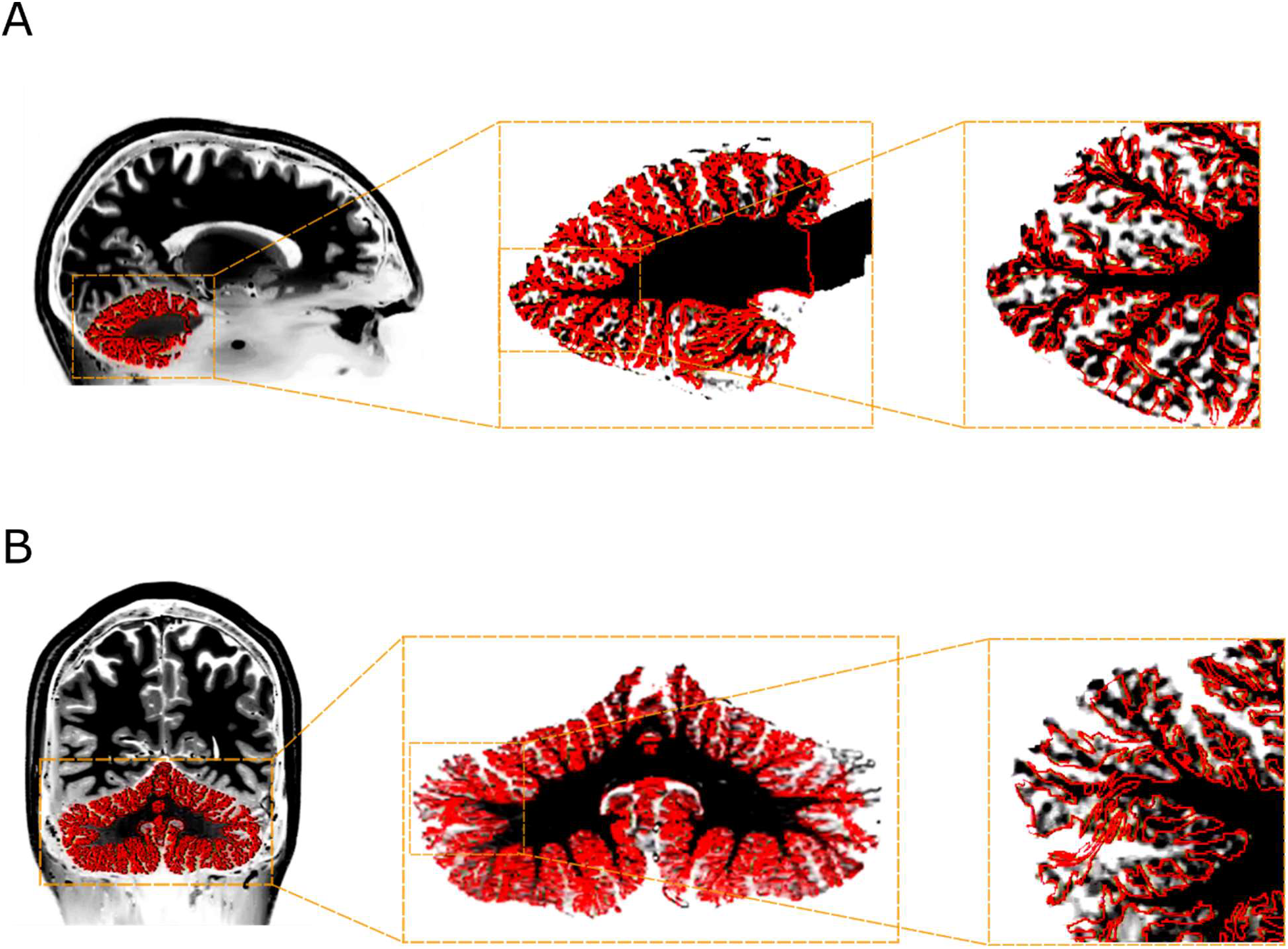
High-resolution (0.25 mm isotropic) MRI volumetric data superposed on the reconstructed cerebellar cortex based on down-sampled (1 mm isotropic) data (red). Sagittal (A) and coronal (B) cross-sections. The images have been contrast-enhanced to better visualize the cerebellar boundaries.

## Discussion

A novel technique for reconstructing the cerebellar cortex was presented. The benefit of this method with respect to an original reconstruction as in Sereno et al. (2020), is that it requires only standard-resolution MRI data. Sereno et al. (2020) showed by empirical testing that a resolution of <0.2 mm is needed to properly resolve the cerebellar folia. This number can also be achieved by a back-of-the-envelope estimation: cerebellar volume is about 15% of the cerebral volume (Hoogendam et al., 2012) while the cerebellar cortical area is 80% of the cerebral cortical area, meaning that the cortical surface area has to be packed about five times more densely in the cerebellum, necessitating a five times higher isotropic resolution. Since the typical MRI resolution required to properly resolve the cerebral cortex is 1 mm isotropic resolution, we would expect the resolution to resolve the folia to be around 0.2 mm. Because this resolution is only practically attainable in high-field ex-vivo MRI scans, this way of reconstructing the cortex directly from the volumetric data is unfeasible for in-vivo scans using contemporary MRI technology. 1 mm^3^ resolution, however, can be achieved in-vivo with widely used commercial MRI scanners, meaning that this technique enables approximate reconstruction of the cerebellar cortex in-vivo and without using specialized hardware. Critically, this means that data that have already been collected with standard scanning hardware and protocols can be reexamined and the cerebellar cortex approximately reconstructed using this technique.

The reconstruction technique presented here results not in a reconstruction with the quality as that which was presented in Sereno et al. (2020), or FreeSurfer’s reconstruction of the cerebral cortex, as is clear from superposing the surface reconstruction on the volumetric data. A more careful analysis of the limitations of this technique is the next step and will have to be completed when more high-resolution surface reconstructions of the cerebellar cortex are available. Currently, the only veridical reconstruction of the cerebellar cortex to the authors’ knowledge is the surface template used in the current study which naturally cannot be tested to itself. When more validation data are available and a suitable goodness-of-fit measure can be attained, the method can also be further refined. While the reconstruction technique presented here has limitations, particularly at the folia-level spatial scale, it is clear that it is considerably better than the current standard FreeSurfer tessellation (Figure 3) and the most accurate approximation of the true cerebellar cortex that can be achieved with standard resolution MRI data to date.

The reconstruction is generally better in the posterior part of the cerebellum and the more medial parts. Comparing the reconstruction results with higher-resolution data (Figure 4), we could clearly see that the overall placement and convex hull of the cerebellum are correct, as are most lobules. However, the folia were often not captured in a correct way. The spatial scale where this technique breaks down is thus between the lobular and foliar level. A prominent question is whether the variant performance in different cerebellar regions is due to limitations in the numerical technique or a higher degree of subject variance in the anterior lobe of the cerebellum. Typically, cortical areas engaged in higher order functions are associated with a higher degree of inter-subject variance (Mueller et al., 2013; Ren et al., 2020). Since the anterior lobe of the cerebellum is mainly associated with motor functions and the posterior lobe with cognitive functions, the reverse would be true for the cerebellum if the differential performance of the reconstruction in these regions are indeed due to inter-subject variance and not for purely technical reasons. A further investigation of this is warranted when more validation data are available, and a quantitative goodness-of-fit measure can be calculated as a function of spatial region.

## Conclusion

A new technique for reconstructing the cerebellar cortex from standard resolution MRI data was presented. Because the technique works by morphing a high-resolution surface template, topographic data can easily be mapped between the template and the subject-specific reconstruction, facilitating population studies and visualizations, e.g., on inflated or flat map surfaces. While the technique has limitations and needs to be validated further, it is the most veridical representation of the cerebellar cortex attainable with standard MRI to date.

## Acknowledgements

The authors would like to extend their gratitude to Dr Martin Sereno who provided the high-resolution cerebellar template data, Dr Koen Van Leemput and Dr William (Sandy) Wells who helped guiding the technical work and Dr Padmavathi (Sundaram) Patel for consultation.

This research was funded by the NIH Neuroimaging Training Program (NTP) grant 5T32EB001680 and NIH-funded P41 Center for Mesoscale Mapping (1P41EB030006). The content is solely the responsibility of the authors and does not necessarily represent the official views of the National Institutes of Health. The authors declare that there is no conflict of interest regarding the publication of this article.

